# A compendium of novel genomics technologies provides a chromosome-scale assembly and insights into the sex determining system of the Greenland Halibut

**DOI:** 10.1101/2021.06.18.449053

**Authors:** A-L Ferchaud, C Mérot, E Normandeau, I Ragoussis, C Babin, H Djambazian, P Bérubé, C Audet, M Treble, W Walkusz, L Bernatchez

## Abstract

Despite the commercial importance of Greenland Halibut (*Reinhardtius hippoglossoides)*, important gaps still persist in our knowledge of this species, including its reproductive biology and sex determination mechanism. In this study, we combined single molecule sequencing of long reads (Pacific Sciences) with Chromatin Conformation Capture sequencing (Hi-C) data to provide the first chromosome-level genome reference for this species. The high-quality assembly encompassed more than 598 Megabases (Mb) assigned to 1 594 scaffolds (scaffold N50 = 25 Mb) with 96 % of its total length distributed among 24 chromosomes. The investigation of its syntenic relationships with other economically important flatfish species revealed a high conservation of synteny blocks among members of this phylogenetic clade. Sex determination analysis revealed that flatfishes do not escape the rule applied to other teleost fish and exhibit a high level of plasticity and turnover in sex-determination mechanisms. A whole-genome sequence analysis of 198 individuals allowed us to draw a full picture of the molecular sex determination (SD) system for Greenland Halibut, revealing that this species possesses a very nascent male heterogametic XY system, with a putative major effect of the sox2 gene, also described as the main SD driver in two other flatfishes. Interestingly, our study also suggested for the first time in flatfishes that a putative Y-autosomal fusion could be associated with a reduction of recombination typical of early steps of sex chromosome evolution.

## INTRODUCTION

The comprehension of sex-determining (SD) mechanisms is of high importance in biology, as they affect developmental processes, the evolution of genomes, and are responsible for determining sex ratio in a population, a key demographic parameter for its viability and stability (Bull 1983, Uller et al 2007). In recent years, genetic analyses in non-model species have revealed a greater degree of sex chromosome diversity and turnover than previously appreciated (Abbott et al 2017). In fact, some lineages, including lizards, fish, amphibians, insects, and plants show frequent changes in the location of SD genes and high rates of turnover of sex chromosomes. This led Furman et al (2020) to propose a new conceptual framework, in which sex chromosome evolution is more cyclical than linear and in which exceptions are the rules in sex determination (McGrath 2020). In this cycle, a new master sex-determining locus arises on an autosome, leading to sex chromosome formation generally characterized by recombination suppression between the Y and X, or the Z and W. As under a classic sex chromosome evolution model, this may progress towards increased divergence between male and female chromosomes, up to chromosome heteromorphism characterized by genomic degradation including rearrangements and loss of genetic material (Charlesworth et al 2005; Bachtrog 2013). However, sex chromosomes and sex-determining regions can also undergo turnover with either the evolution of a new sex-determining gene or the transposition of the sex-determining gene to a new location in the genome, leading to another cycle (Bachtrog 2006). The increasing availability of high-quality genome assemblies in non-model taxonomic groups is now providing exciting opportunities to revisit the evolution of genetic sex-determination.

Fish form a particularly relevant group to investigate the evolutionary dynamics of sex chromosomes, as they exhibit a remarkable gonad development plasticity and a high evolutionary SD turnover (Kitano and Peichel 2012). Fish display highly diverse chromosome SD systems, including models analogous to the XX/XY (male heterogametic) and ZZ/ZW (female heterogametic) systems of mammals and birds respectively, but also others involving multiple sex chromosomes (Capel 2017). In accordance with this rapid turnover, chromosome heteromorphisms are rare in fish, which generally show limited sex chromosome differentiation (Chen et al 2014), characteristic of early sex-determination mechanisms. Until recently, the DM-domain gene on the Y chromosome (dmy) gene was the only known gene coding for a functional protein in the Y-specific chromosome region (Nanda et al 2002, Kondo et al 2006) and was described as the master SD gene in a wide range of invertebrates and vertebrates (including fish). However, thanks to new genomics tools, the genomic features underlying SD have been more intensively studied and several other strong candidates of master SD genes have been found in fish (amhy, gsdf, amhr2, sdY, Guerrero-Estévez and Moreno-Mendoza 2010, Yano et al 2013). These novel master SD genes highlight the importance of non-transcriptional factors in sex determination and differentiation. In fact, three of these genes (amhy, amhr2, and gsdf) are involved in the transforming growth factor beta (TGF-β) signaling pathway, while the sdY, dmrt1, and dmy genes code for transcription factors (Rastetter et al 2015). Insights into SD genomics goes beyond master genes in fish, as recent studies have documented mixed determination involving both genetic and environmental factors, polygenic SD systems, extensive within-species variation in SD system, and complex rearrangements of sex chromosomes favouring species diversification (Capel 2017, Martinez et al 2014). Yet, with such dynamism and variability in fish sex chromosome evolution, we need improved genomic resources in non-model species, including chromosome-scale reference genomes and whole-genome sequences from both sexes.

In exploited species, improved genomic resources and the understanding of the genetic basis for SD is also relevant to manage fisheries in order to meet the need for seafood around the world, and to secure the economies and livelihoods that depend on them (Bernatchez et al 2017). Genomics approaches allow for the investigation of genome wide diversity and to accurately identify fish stock within commercial species (Ovenden et al 2013). Moreover, accounting for sex-differentiation and sex-linked loci is necessary to avoid unexpected biases due to sex-associated markers (Benestan et al 2017) as well as to improve the management of sex-ratio. This is particularly true for species harbouring a size-related sexual dimorphism like flatfish species, generally favouring females in their size dimorphism (e.g. European Plaice (*Pleuronectes platessa*; Rijnsdorp and Ibelings 1989), Turbot (*Scotphtalmus maximus*; Imsland et al 1997), Atlantic Halibut (*Hippoglossus hippoglossus*; Hagen et al 2006). In such species, sex control and/or identification is of high importance either for flatfish production in aquaculture or sustainable fisheries. The selection of large individuals in fisheries is a well-established concept in fisheries management that increases yield per recruit. However, in the case of sex-related sexual dimorphism, fisheries targeting large individuals may reduce the stock’s reproductive potential, produce bias in the assessment of the number of reproductive females, and feed erroneous data to fisheries management decision makers (Keyl et al 2015). Overall, applied and fundamental evolutionary biology thus benefits from moving beyond genetic markers towards contiguous reference genomes annotated for functional elements and documented for sex-determination.

Greenland Halibut (*Reinhardtius hippoglossoides*) is one of the main demersal fish exploited by fisheries in Eastern Canada (Gulf of St. Lawrence, Newfoundland and the Arctic), representing around 4 000 tones of landings per year across the two last decades (DFO, 2018). However, it suffers from a lack of genomic resources, since no karyotype, linkage map nor genome assembly has been developed so far, thus hampering the use of genomics for improved management. Since juvenile growth potential and size are sexually dimorphic in this species (Ghinter et al 2019), it is valuable to better understand the sex-determining mechanisms to control sex imbalance in catches. Moreover, the recent genome assemblies of flatfish species (Figure 1, Table1, Chen et al 2014, Figueras et al 2016, Shao et al 2017, Einfeldt et al 2021) together with studies on sex-determining loci in this group (Chen et al 2014, Liang et al 2018, Drinan et al 2018, Einfeldt et al 2021) provides an opportunity to investigate the evolutionary turnover of sex chromosomes and sex-determination systems. Flatfishes studied so far are characterized by a variable heterogametic system and different sex-determining genes. For instance, a similar female heterogametic system (ZZ/ZW) is used both by the Tongue Sole (*Cynoglossus semilaevis*) and the Turbot, but the candidate SD drivers differ, being sox2 in the latter (Martinez et al 2021, Table1), and dmrt1 in the former species (Chen et al 2014). Japanese Flounder, *Paralichthys olivaceus* (also called the Olive flounder,) has an XX/XY system (Liang et al 2018) but no sex determination gene has been found (Shao et al 2017) and environment is known to alter the gender (Ospina-Alvarez and Piferrer 2008). Within the righteye flounder family (*Pleuronectidae*), the *Hippoglossinae* subfamiliy harbors variable genetic SD systems. The Pacific Halibut (*Hippoglossus stenolepis*) has a ZZ/ZW system and several sex-linked loci aligned in three different linkage groups (Drinan, Loher, & Hauser 2018) while its sister-species, the Atlantic Halibut (*H. hippoglossus*), has a XX/XY system and a putative SD factor called gsdf on chromosome 12 (Palaiokostas et al. 2013; Einfeldt et al 2021).

**Figure 1:**
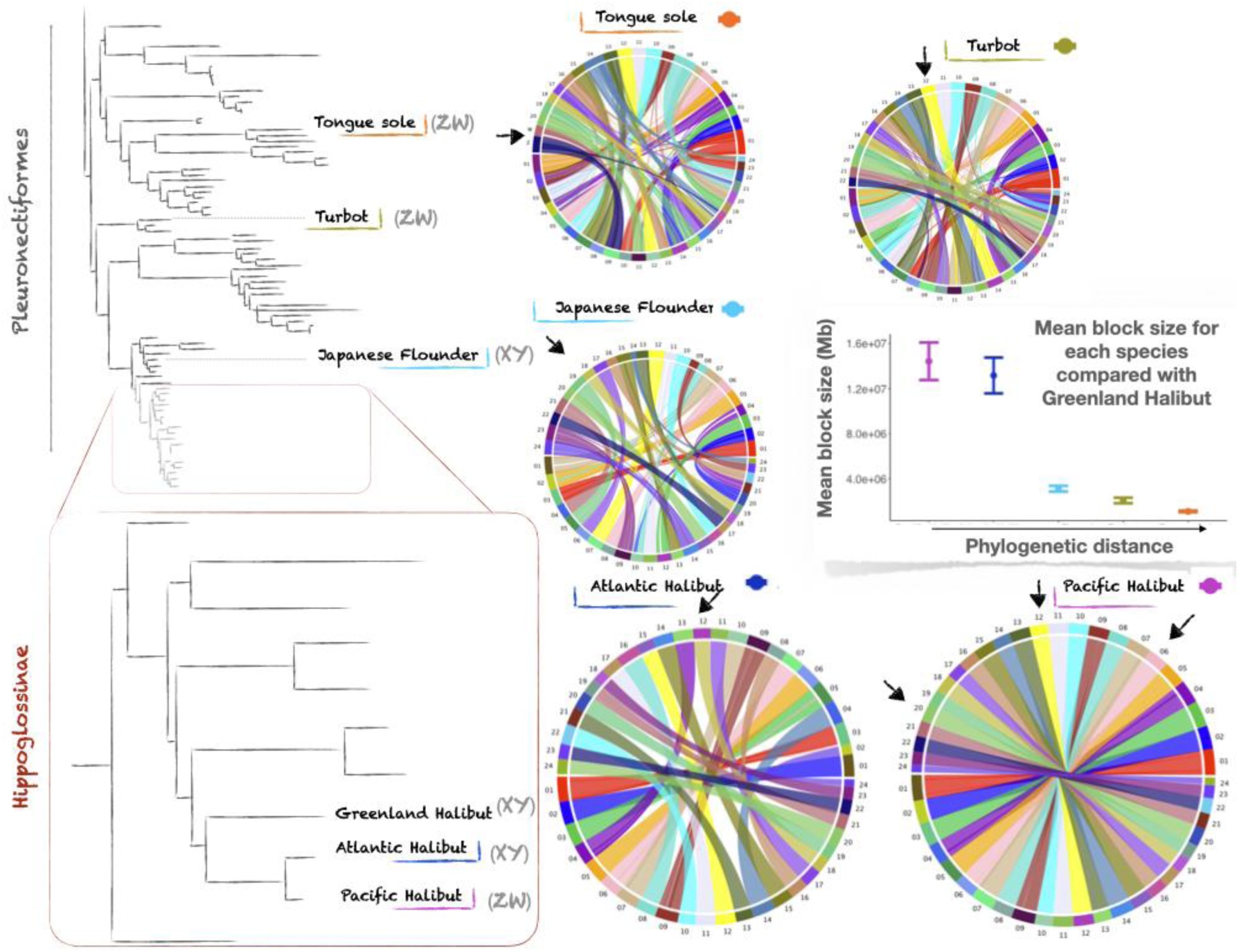
Phylogenetic relationships and syntenic distances between important flatfish species in fisheries and aquaculture. Phylogenetic trees were retraced according to previous studies, Betancour et al (2014) for the top one and Vinnikov et al (2018) for the zoom into the sub-family *Hippoglossinae* framed in red. Only the names of relevant species compared in our studies are reported as well as the SD system, either “XY” for a male heterogametic system (XX/XY) or “ZW” for a female heterogametic system (ZZ/ZW). The XY system for Greenland Halibut has been assessed in the present study. Circos plots showing synteny between the Greenland Halibut (displayed in low half circle) and the Tongue Sole, Turbot, Japenese Flounder, Atlantic and Pacific Halibut. Black arrows in each circos plot refers to the sex chrosomosomes or the location of sex-related markers. A positive relationship between the mean block size and phylogenetic distance is also reported. Colors used for reporting the standard deviation and the mean of block size are indicated next to each species name respectively. Note that the phylogenetic distance reported on the x-axis is qualitative only and that species have been positioned from the most closely related species to the most distant species. The same color code as the one near the circus plots is used.

**Table 1:** Statistics of genome assemblies of the important flatfish species in fisheries and aquaculture. n.a. is used for data not available ^(1)^ This study, ^(2)^ Drinan, Loher, & Hauser, 2018, ^(3)^ Einfeldt et al 2021, ^(4)^ Liang et al 2018, ^(5)^ Martinez et al 2021, ^(6)^Cheng et al 2014

|  | Greenland Halibut | Pacific Halibut | Atlantic Halibut | Japanese flounder | Turbot | Tongue sole |
| --- | --- | --- | --- | --- | --- | --- |
| Species latin name | *Reinhardtius*<br>*hippoglossoides* | *Hippoglossus*<br>*stenolepis* | *Hippoglossus*<br>*hippoglossus* | *Paralichthys*<br>*olivaceus* | *Scophthalmus*<br>*maximus* | *Cynoglossus*<br>*semilaevis* |
| total sequence length | 598,525,902 | 594,145,200 | 596,792,615 | 545,775,252 | 524,979,463 | 470,199,494 |
| number of scaffolds | 1,594 | 120 | 57 | 9,525 | 22 | 31,181 |
| max scaffolds lengh | 31,982,681 | 32,413,155 | 30,753,830 | 27,810,759 | 31,901,587 | 34,528,841 |
| scaffold N50 | 25,015,410 | 24,985,257 | 26,312,018 | 3,817,360 | 24,811,384 | 509,861 |
| scaffold L50 | 11 | 11 | 11 | 40 | 10 | 289 |
| total number of<br>chromosomes | 24 | 24 | 24 | 24 | 22 | 21 |
| Genbank accession number | GCA_006182925.3 | GCA_013339905.1 | GCA_009819705.1 | GCA_001904815.2 | GCA_003186165.1 | GCA_000523025.1 |
| Reference | This study | International | Einfeldt et al 2021 | Shao et al 2017 | Martinez et al | Chen et al 2014 |
|  |  | Pacific Halibut<br>Commission |  |  | 2021 |  |
| **Name of the assembly** | UL_REHI_2.0 | IPHC_HiSten_1.0 | fHipHip1 | ParOli_1.1 | ASM318616v1 | Cse_v1.0 |
| **Complete BUSCO** | 98.04% | na | 96.90% | na | na | na |
| **Repeat elements** | 8.20% | 7.28% | 7.14% | 4.92% | 6.35% | 6.08% |
| **Genetic sex-determinism** | XX//XY | ZZ/ZW | XX/XY | XX/XY | ZZ/ZW | ZZ/ZW |
| **SD genes or putative SD genes** | sox2, sox9a2, gdf6<br>(1) | no gene has been<br>specifically<br>identified (2) | gsdf (3) | Cyp11a (4) | sox2 (5) | dmrt1 (6) |
| **Homolog of sex**<br>**chromosome in Greenland**<br>**Halibut** | - | GH06, GH12, G20 | GH18 | GH19 | GH14 | GH09 |

In this context, the first goal of this study was to provide high-quality genomic resources and to investigate the genetic basis of sex determination in Greenland Halibut. We produced the first chromosome-scale scaffolded genome assembly for this species, combining single molecule sequencing of long reads (Pacific Sciences) with Chromatin Conformation Capture sequencing (Hi-C) data. Furthermore, we also identified and characterized the molecular SD system by mapping whole genome sequencing data of hundreds of individuals onto the assembly to screen for divergent genetic markers between sexes. Finally, we analyzed our results in a comparative genomic framework by assessing synteny (i.e. block of genes in the same relative positions in the genome) with related species of flatfish and targeting candidate sex-determining genes in this group.

## METHODS

### Reference genome assembly

We built a reference genome assembly for the Greenland Halibut through two steps. First, we created a draft assembly using PacBio sequencing approach (Pacific Biosciences, Menlo Park, California) and then we used Dovetail™ Hi-C (Lieberman-Aiden et al 2009) – from Dovetail Genomics to increase contiguity, break up mis-joins, and orient and join scaffolds into chromosomes.

### Sampling

The individual used for the reference was a juvenile female caught in the St. Lawrence River Estuary (48°39′11″N, 68°28′37″W) with a Comando-type trawl aboard the CCGS Leim (Fisheries and Oceans Canada [DFO] survey) at the end of May 2016 and kept alive in common garden environment at the Maurice Lamontagne Institute (Ghinter et al 2019). The female was anaesthetized for 5 min in a solution of MS 222 (tricaine methane sulfonate 0.18 g L^-1^; Sigma - Aldrich Co, Missouri, USA). Blood was then sampled from the caudal artery using a 23-gauge needle in a 1ml TB syringe (Becton, Dickinson & co., New Jersey, USA), both of which were previously heparinized in a 100 U ml ^-1^ heparin solution. Blood was immediately flash frozen in liquid nitrogen and then stored at -80°C until further manipulation, as well as multiple tissues including liver, muscle, heart, brain, eye, gonads, gall bladder and kidney that were excised from the fish.

### DNA extraction, sequencing and assembly

High Molecular Weight DNA was extracted using Qiagen MagAttract kit, (Valencia, CA). Then a library was created using the SMRTbell Template Prep Kit 1.0 (Pacific Biosciences, Menlo Park, California), which includes a DNA Damage Repair step after size selection. Size selection was made for >10 kb using a Blue Pippin instrument (Sage Science, Beverly, Massachusetts) according to the manufacturer recommended protocol for 20 kb template preparation. Five µg of concentrated DNA was then used as input for the library preparation reaction. Library quality and quantity were assessed using the Pippin Pulse field inversion gel electrophoresis system (Sage Science), as well as with the dsDNA Broad Range Assay kit and Qubit Fluorometer (Thermo Fisher). Single-Molecule Real Time sequencing was performed on the Pacific Biosciences RS II instrument at McGill University using an on-plate concentration of 125 pM, P6-C4 sequencing chemistry, with magnetic bead loading, and 360-minute movies. A total of 14 SMRT Cells were run, generating a mean sequence coverage of ∼70-fold, with half of the 522 Megabases (Mb) of data contained in reads longer than 10.6 kb. An initial assembly was carried out on the PacBio long reads using Wtdbg v2.5 (-x sq gg 0.7g; Ruan and Li (2020), resulting in 2 037 contigs (N50 = 31.7 Mb), for a total assembly size of 598 MB. One Hi-C proximity ligation library was generated from a flash frozen piece of muscle of the same individual using the Dovetail’s ™ Hi-C Kit v.1.0 to aid with assembly scaffolding. This kit employs the restriction enzyme *Dpn*II which recognizes the sequence 5’ /GATC 3’. The long-range library was then sent to Dovetail Genomics to be sequenced and run with the initial PacBio assembly through Dovetail’s HiRiseTM scaffolding pipeline. The completeness of the final assembly was assessed with BUSCO v 4.0.5 (Simão et al 2015), the eukaryota_odb10 database (created in October 2020) and the -m genome option. Occurrence of repeat elements in our assembly as well as in Atlantic and Pacific halibut was assessed using RepeatMasker v.4.0.6 (Smit, Hubley and Green, 2013).

### Synteny with other flatfish species

Synteny blocks between the genomes of Greenland Halibut and other flatfish species for which a reference genome is available (Tongue Sole, Turbot and Japanese Flounder as well as Atlantic and Pacific Halibut) were computed using SyMAP v4.2 (Soderlund, Bomhoff and Nelson 2011). Genomic sequences were first aligned using promer/MUMmer (Kurtz et al 2004). Raw anchors resulting from MUMmer were clustered into (putative) gene anchors, filtered using a reciprocal top-2 filter and used as input to the synteny algorithm (Soderlund et al 2006). The algorithm constructs maximal-scoring anchor chains based on a given gap penalty. It also searches a range of gap penalties to generate the longest chains subject to several quality criteria, which are based on the Pearson correlation coefficient applied to the anchors in the chain as well as the anchors in its bounding box. The chains are not required to be entirely colinear and may incorporate local inversions relative to the overall chain orientation. Synteny analysis results were visualized using the Chromosome Explorer and the Circle view options in SyMAP v4.2. Finally, we named scaffolds by mapping our assembly to the Pacific Halibut assembly (IPHC_HiSten_1.0, Genbank accession: GCA_013339905.1).

### Transcriptome assembly and annotation of the assembly

In order to annotate the reference assembly, total RNA from 8 different tissues (liver, muscle, heart, brain, eye, gonads, gall bladder and kidney) from the same individual was extracted using the RNeasy Mini kit from Quiagen then treated using DNase I, Amplification Grade (1 unit per μl; Invitrogen, Carlsbad, CA, USA) following the manufacturer’s protocol. RNA concentration of each tissue was quantified with QuantiT Picogreen dsDNA Assay Kit 669 (Invitrogen) and fragment size distribution was estimated with an Agilent BioAnalyzer before being shipped to the Genome Québec Centre d’Expertise et de Services (Montréal, Canada) to prepare libraries and perform sequencing. One RNA-seq library was constructed for each tissue using the **NEBNext**^**®**^ **Ultra**^**™**^ **|| Directional RNA Library Prep Ki** (Illumina, San Diego, Calif., USA) and sequenced with a 2 × 100bp (paired-end) read module using the NovaSeq 6000, generating 50M reads.

For each of the 8 paired-end tissue-specific RNAseq datasets, raw reads were trimmed with trimmomatic (v0.36, ILLUMINACLIP:univec.fasta:2:20:7, LEADING:20, TRAILING:20, SLIDINGWINDOW:30:30, MINLEN:80). The tissue-specific paired-end datasets were then each assembled separately using Trinity (v2.1.1, --seqType fq, --min_contig_length 200) to produce 8 assembled transcriptomes that contained between 68 481 and 276 168, with a mean of 131 038, assembled expressed sequences. The 1 048 307 assembled expressed sequences were used to annotate the assembled genome using the GAWN pipeline v0.3.4 (https://github.com/enormandeau/gawn). The annotations were then simplified by using the included simplify_genome_annotation_table.py script, which removes consecutive almost-identical annotations from the genome annotation table.

### Genomic sex determination

#### Sampling of individuals, DNA extraction, libraries and sequencing

A total of 432 Greenland Halibut were collected aboard the R/V Paamiut, from October to November 2016, as part of the annual multispecies survey conducted by the Department of Fisheries and Oceans in Northwest Atlantic Fisheries Organization Subarea 0 in Davis Strait, Canada. Each fish collected has been measured and weighed as well as phenotypically sex identified. From those individuals, 99 females and 99 males have been selected for sex determination analysis (see sample selection details in Supplementary Material).

Genomic DNA was extracted using a salt-extraction protocol (Aljanabi and Martinez 1997) with a RNase A treatment 645 (Quiagen). DNA quality of each extract was evaluated with nanodrop and on a 1% agarose gel electrophoresis. Following Therkildsen and Palumbi 2017, we removed DNA fragments shorter than 1kb by treating each extract with Axygen magnetic beads in a 0.4:1 ratio, and eluted the DNA in 10mM Tris-Cl, pH 8.5. We measured DNA concentrations with Accuclear ultra high sensitivity dsDNA quantification kit (Biotium) and normalized all samples at a concentration of 5ng/ μL. Then, sample DNA extracts were randomized, distributed in plates (96 -well see details about randomization and number of plates used in Supplementary Material) and re-normalized at 2ng/μL.

Whole-genome high-quality libraries were prepared for each sample according to the protocol described in (Baym et al. 2015; Therkildsen and Palumbi 2016). Briefly, a tagmentation reaction using enzyme from the Nextera kit, which simultaneously fragments the DNA and incorporates partial adapters, was carried out in a 2.5 μl volume with approximately 2 ng of input DNA. Then, we used a two-step PCR procedure with a total of 12 cycles (8+4) to add the remaining Illumina adapter sequence with dual index barcodes and to amplify the libraries. The PCR was conducted with the KAPA Library Amplification Kit and custom primers derived from Nextera XT set of barcodes A, B, C and D (total 384 combinations). Amplification products were purified from primers and size-selected with a two-steps Axygen magnetic beads cleaning protocol, first with a ratio 0.5:1, keeping the supernatant (medium and short DNA fragments), second with a ratio 0.75:1, keeping the beads (medium fragments). Final concentrations of the libraries were quantified with Accuclear ultra high sensitivity dsDNA quantification kit (Biotium) and fragment size distribution was estimated with an Agilent BioAnalyzer for a subset of 10 to 20 samples per plate. Finally, libraries were pooled (see details in Supplementary Material) for sequence lanes of paired-end 150 bp reads on an Illumina HiSeq 4000 at the Norwegian Sequencing Center at the University of Oslo. Given our multiplexing (up to 96 individuals per lane) and genome size (∼ 600 Mb), we targeted to a low coverage sequencing (∼ 1.26 X).

#### Sequencing filtering and processing

Raw reads were trimmed, filtered for quality, mapped to the reference genome (obtained above), cleaned for mapping quality and duplicates reads and then re-aligned using the pipeline available at https://github.com/enormandeau/wgs_sample_preparation, inspired by (Therkildsen and Palumbi 2016) and fully described in Mérot et al (2021). Given the high percentage of the assembly comprised into the anchored 24 chromosomes (> 96 %, see results section), raw reads were mapped against a reduced version of the genome including only the 24 chromosomes and not the unplaced scaffolds.

For low coverage whole genome sequencing (lcWGS) data, the recommended practice is to avoid basing downstream analysis on raw counts of sequenced bases or called genotypes (Nielsen et al 2011), but instead use a probabilistic approach based on the use of Genotype Likelihoods. Several models for computing genotype-likelihood-based on read data have been implemented in the program ANGSD (Korneliussen, Albrechtsen and Nielsen et al 2014), which is currently the most widely used and versatile software package for the analysis of lcWGS (Lou et al 2021). Therefore, the program ANGSD v0.931 was used for most of our subsequent analyses, according to the pipeline documentation available at https://github.com/clairemerot/angsd_pipeline. For all analyses, input reads were filtered to remove reads with a samtools flag above 255 (not primary, failure and duplicate reads, tag - remove_bads = 1), with mapping quality below 30 (-minMapQ 30) and to remove bases with quality below 20 (-minQ 20). We also filtered in order to keep only SNPs covered by at least one read in at least 30 % of individuals (-minInd 60) and remove SNPs in putative repeated regions allowing a maximum depth of 3 times the number of individuals (-setMaxDepth 594). Finally, for most of the subsequent analysis (unless mentioned otherwise) we kept SNPs with minor allele frequency above 5%.

First, we ran ANGSD to estimate genotype likelihoods (GL) with the GATK model (-doGlf 2 -GL 2 -doCounts 1), the spectrum of allele frequency (-doSaf 1) and minor allele frequency (-doMaf The major allele was based on the genotype likelihood and was the most frequent allele (-doMajorMinor 1). From this first analysis, we generated (i) a beagle file with GL estimates and (ii) a list of variants passing those filters and their respective major and minor alleles that were used for most subsequent analysis. The R program (R Core Team 2020) was employed for graphic output in subsequent analyses, *via* the package ggplot (Wickham 2016).

#### Genome-wide coverage difference between sexes

Total depth along the entire genome was calculated using the function -doDepth 1 in ANGSD, excluding high coverage (-setMaxDepth 594 and -maxDepth 1000) and low coverage regions (-minInd 60), along with the following arguments for depth: -dumpCounts 2 -doCounts 1 and then averaged across sliding-windows of 25Kb with a step of 5 Kb using the winScan() function of the R package windowscanr (Tavares 2021). The sliding approach allowed us to visualize individual coverage along the genome, while a ratio of total depth of females above total depth of males was also plotted using ggplot(). This ratio is expected to deviate from 1 in genomic regions in which a sex-bias in coverage exists.

#### Genome wide variation between sexes

Genome-wide variation across samples was explored using PCAangsd (Meisner and Albrechtsen 2018) on the genotype likelihoods. This program extracts a covariance matrix that is then decomposed into principal component analysis (PCA) with R using a scaling 2 transformation adding an eigenvalues correction, to obtain the individuals PC scores (Legendre and Legendre 1998).

#### Identify genomic regions implied in sex determination

In order to identify the genomic regions involved in the sex differentiation, two analyses were performed. First, we ran PCAangsd on genotype likelihoods in non-overlapping windows of 1 000 SNPs in each chromosome to extract local covariance matrices and obtained local PCAs of genomic variation (as detailed above). For each local PCA, we analyzed the correlation between individuals PC1 scores and PC scores from the global PCA performed on the entire genome. The resulting correlation indices were then plotted across their relative position in each chromosome. Second, genome-wide Fst was estimated between males and females. To do this, allele frequency spectrum (-doSaf 1) and minor allele frequencies were calculated for each sex with the previous list of variant positions (-sites) and their polarisation as major or minor alleles (-doMajorMinor 3). Genome-wide Fst was estimated using the realSFS function in ANGSD between males and females and then summarized across sliding-windows of 25 Kb with a step of 5 Kb.

Observed heterozygosity was computed for each sex and each SNP using the function - doHWE in ANGSD and then averaged across sliding-windows of 25Kb with a step of 5 Kb using the winScan() function of the R package windowscanr.

#### LD estimation

For the two chromosomes associated with sex determination (see results below), intra-chromosomal as well as inter-chromosomal linkage disequilibrium was calculated within each sex for a reduced number of SNPs, filtered with more stringent criteria (MAF > 10%, at least one read in 75 % of the samples). Pairwise R^2^ were calculated using ngsLD (Fox et al 2019) based on genotype likelihood reported in the beagle file obtained previously. The 10th percentile of pairwise R^2^ values was averaged across windows of 250 Kb and plotted for each sex using custom scripts available at https://github.com/clairemerot/angsd_pipeline. Then changes in LD along the sex identified chromosomes were analyzed by calculating mean R^2^ for males and females separately.

#### Mapping known sex locus

To localize homolog genes of known sex-associated genes in our assembly, we aligned all sex-associated markers from Drinan et al (2018) as well as others previously described to have a sex-determination effect in flatfish or teleost species, to our reference assembly using LAST v.1179 (Kielbasa et al 2011; see Table S1 for details about those genes and their accession number).

## DATA AVAILABILITY

Genome assembly is available upon the GCA_006182925.2 accession. Raw data for the 8 RNA-seq lanes are available on NCBI upon following accession numbers: SAMN18221283 to SMAN18221290 and 198 individual whole genome sequences are submitted under the BioProject ID PRJNA737354 on NCBI SRA and will be available upon publication.

## RESULTS AND DISCUSSION

### A high-quality reference genome serving flatfish management and research

We generated a final genome assembly of 598.5 Mb, with overall high contiguity (N50=25MB, N90=20MB), of which 96.35% was comprised into 24 chromosome-length scaffolds (GenBank accession: GCA_006182925.3). These 24 scaffolds are consistent with the number of diploid chromosomes (2n=48) for Atlantic Halibut (Einfeldt et al 2021) and Pacific Halibut (GCA_013339905.1). The Hi-C data supported a high degree of accuracy in the overall assembly into these 24 scaffolds, as indicated by the strong concentration of data points along the diagonal rather than elsewhere in the contact maps (Figure S1). Altogether, this suggests that these 24 scaffolds can be considered as chromosomes. The completeness of the assembly based on single-copy orthologs across eukaryotes was 98.04%, with 96.47% complete and single copy orthologs and 1.57% duplication.

There were 60 313 transcripts detected from multi-tissue transcriptomics data, with 97.7% of them (58 908 transcripts) within the 24 anchored chromosomes. A total of 19 252 genes and pseudogenes were annotated, of which 19 053 (99%) were located within the 24 chromosomes. Repeated elements accounted for 8.20% of chromosome length (49.1 Mb). These measures of completeness and assessments of repeat elements in Greenland Halibut were highly concordant with the statistics obtained from Atlantic and Pacific Halibut as well as the three other flatfish species with reference genome and important for fisheries and/or aquaculture production (Table 1). SyMap revealed shared syntenic blocks with other flatfish species, allowing the identification of homologous chromosomes (Table 2 and Figure 1). Most of the chromosomes harbor a single syntenic block compared with the most closely-related *Hippoglossinae*, the Pacific Halibut (16 out of 24 chromosomes) and the Atlantic Halibut (15/24). In contrast, mapping with the three other flatfish species displayed several syntenic blocks for all chromosomes (Table 2). As a result, syntenic blocks are larger with Pacific and Atlantic Halibut than with the three other flatfishes (Table 2, and Figure 1). Figure 1 displays this negative relationship between the size of syntenic blocks and phylogenetic distance.

**Table 2:**
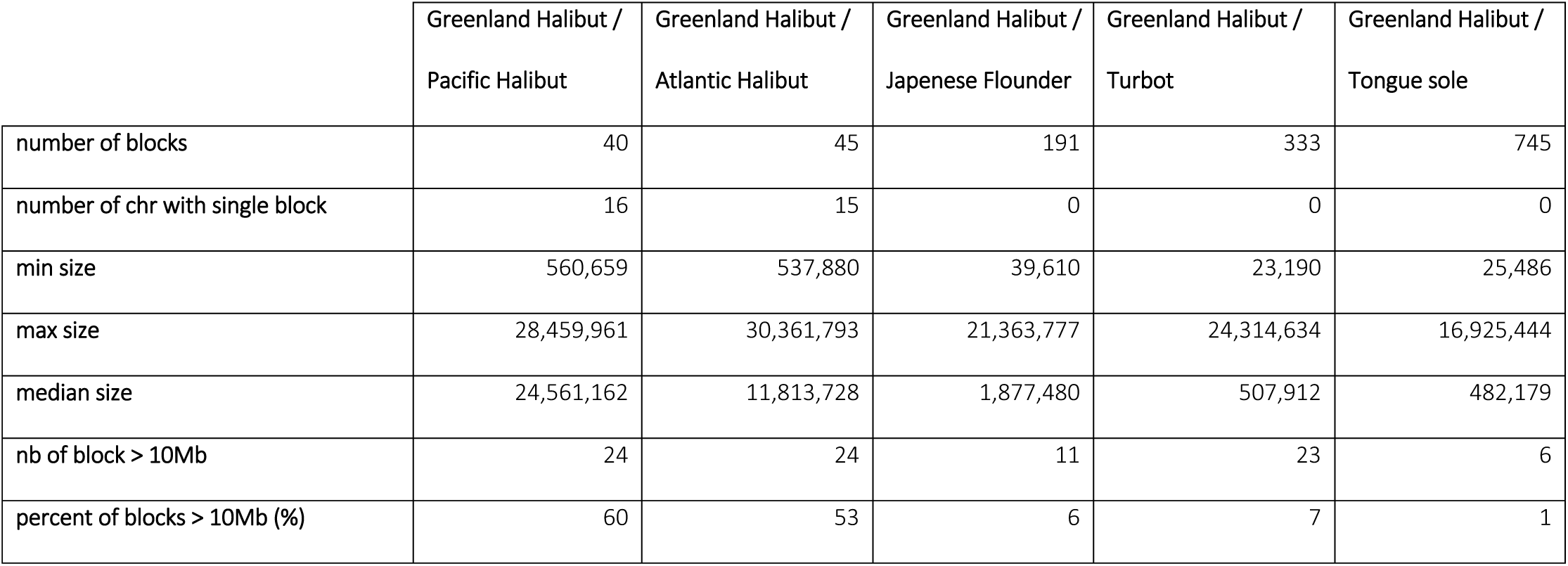
Syntenic relations between Greenland Halibut and related commercial flatfish species.

|  | Greenland Halibut /<br>Pacific Halibut | Greenland Halibut /<br>Atlantic Halibut | Greenland Halibut /<br>Japanese Flounder | Greenland Halibut /<br>Turbot | Greenland Halibut /<br>Tongue sole |
| --- | --- | --- | --- | --- | --- |
| number of blocks | 40 | 45 | 191 | 333 | 745 |
| number of chr with single block | 16 | 15 | 0 | 0 | 0 |
| min size | 560,659 | 537,880 | 39,610 | 23,190 | 25,486 |
| max size | 28,459,961 | 30,361,793 | 21,363,777 | 24,314,634 | 16,925,444 |
| median size | 24,561,162 | 11,813,728 | 1,877,480 | 507,912 | 482,179 |
| nb of block > 10Mb | 24 | 24 | 11 | 23 | 6 |
| percent of blocks > 10Mb (%) | 60 | 53 | 6 | 7 | 1 |

By generating this chromosome-level genome assembly for Greenland Halibut, among the most contiguous of fish assemblies to date, we provide a genomic resource for this unique species of the genus *Reinhardtius*. This reference genome will now become a fundamental tool to understand the genomic variability of Greenland Halibut in the context of comparative genomics and fisheries applications.

### Nascent XY sex determination system

After mapping the reads of 99 male and 99 female Greenland Halibut samples to the reference genome, cleaning for quality and processing for SNP identification using GL approach, we identified 7 301 470 Single Nucleotide Polymorphisms (SNPs) with a minor allelic frequency (MAF) above 5%. These SNPs were then used for further differentiation analyses.

Genome-wide variation analyzed by a global PCA displayed two clusters corresponding to males and females on PC1, which explained 5.7 % of the genetic variance (Figure S2). Local PCAs on windows of 1 000 SNPs along the genome revealed that this sex differentiation was mainly explained by genetic variation on two chromosomes (Chr10 and Chr21 Figure 2A). The correlation between the 1^st^ PC of the global PCA and each local PCA was very low at the very beginning of each of those two chromosomes but it increased rapidly along both chromosomes. A large portion of each of these two chromosomes explains the total genetic variation observed between the two sexes in PC1. Particularly, the last 66.5 % of Chr10 (16.5 Mb out of 24.8 Mb) and 52.2 % of Chr21 (10.5 Mb out of 20.1 Mb) explained more than 90 % the sex differentiation observed on global PC1 (Figure 2B). Genome-wide Fst estimations between males and females corroborate this result revealing high differentiation on Chr10 and Chr21 (Figure 2C). While the mean genome-wide Fst between the sexes is as low as 0.01, Chr10 and Chr21 harbored a large region with higher Fst values with a mean Fst of 0.21 and 0.15, respectively (Figure 2D). Few localized peaks of sex differentiation were also observed on Chr9, 11, 12, 13 and 15 (Figure 2B).

**Figure 2:**
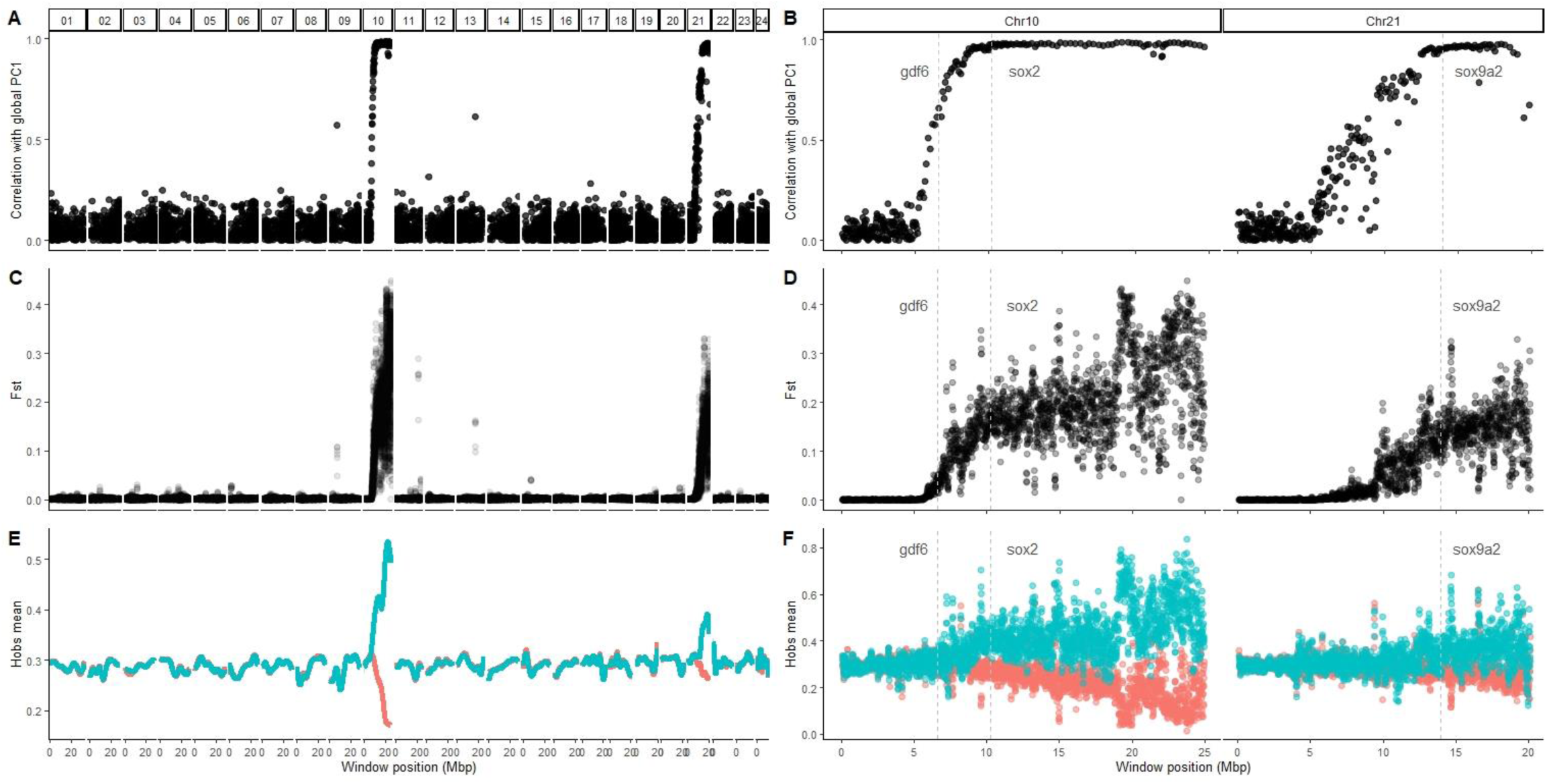
Two genomic regions implied in sex differentiation. **(A)** Correlation between PC1 scores of the local PCAs performed on windows of 1000 SNPs and PC1 scores of the global PCA along their genomic position in each chromosome. **(C)** Fst differentiation estimated between males and females in sliding-windows of 25 Kb within each chromosome. **(E)** Mean observed heterozygosity in each sex smoothed for visualization. **(B), (D)**, and **(F)** are zooms on the two differentiated chromosomes (chr 10 and 21) of the previous values, respectively (A), (C), and (E). Vertical dashed grey lines indicate the positions of detected sex-related candidate genes.

The observed proportion of heterozygotes followed the same pattern, with similar values between males and females across most of the genome (Figure 2E), but with a much higher fraction of heterozygotes in males than females in the two major regions previously described. This difference between males and females was more pronounced in Chr10 than in Chr21. Both Chr10 and Chr21 were also characterized by strong linkage disequilibrium, which was markedly higher in males than in females (Figure 3 and Figure S4), suggesting that recombination events are less frequent in males than females for these genomic regions.

**Figure 3:**
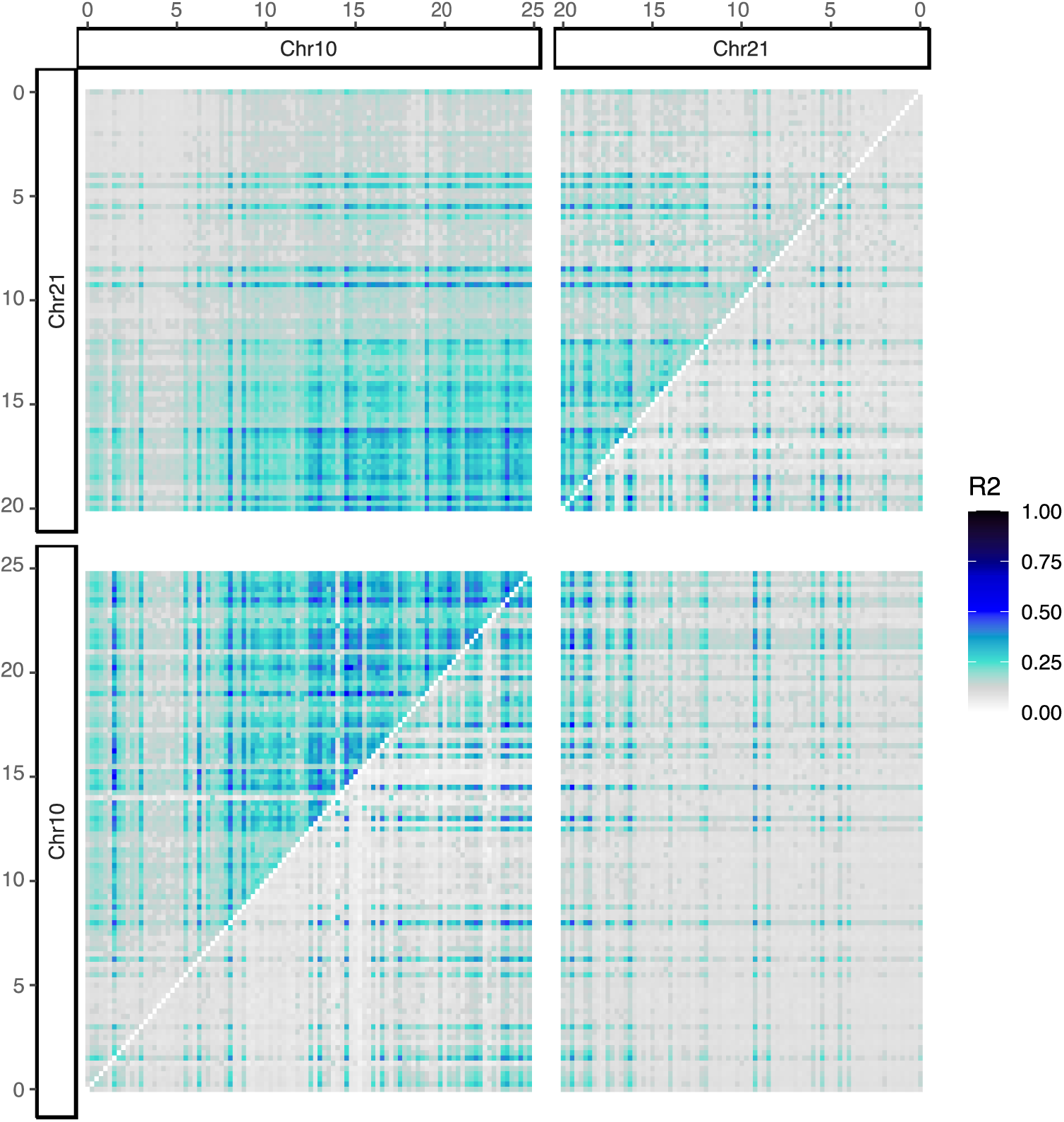
Intra- and Inter-chromosomal Linkage Disequilibrium (LD) among males and females, revealing a putative Y-autosomal fusion. Male values are displayed above the diagonal and female values are displayed below the diagonal. The color scale corresponds to the 10^th^ highest percentile of the R2 value of each SNP pairwise summarized by windows of 250 Kb. Positions are represented in megabases.

In contrast with differentiation and heterozygosity, coverage did not differ between males and females, neither genome-wide nor in regions of sex differentiation (Figure S3), suggesting that Greenland Halibut is not harboring sex chromosome heteromorphism and that the putative sex chromosomes Chr10 and Chr21 have not undergone degradation, which is typical of early SD systems.

Altogether, these results point towards a major role of Chr10 and Chr21 in genetic sex determination. The higher heterozygosity and LD in males suggests that Greenland Halibut has a XX/XY sex determination mechanism. The putative male Y chromosome exhibits two characteristics of sex chromosome evolution; it is divergent from the X chromosome and exhibits a reduction of recombination. However, it does not appear degraded, suggesting very nascent sex chromosomes at an early evolutionary stage. This is corroborated by the syntenic relationships with closely related flatfish species which indicate that homologues of Chr10 and Chr21 are autosome chromosomes. Chr10 and Chr21 neither correspond to locations where sex-determining regions have been defined in other species (Figure 1, Table1). Overall, the resolved sex-determining system in Greenland Halibut is highly consistent with the rapid turnover of sex chromosomes in fish, particularly in flatfish, given that the Atlantic Halibut has also undergone a recent evolution of an XY sex-determination mechanism (Einfeldt et al 2021).

Observing high sex differentiation on two different chromosomes was nevertheless unexpected and puzzling. The Hi-C contact map very strongly supported that Chr10 and Chr21 are distinct chromosomes in the specimen used for the assembly, a juvenile female (Figure S1). Inter-chromosomal linkage disequilibrium was also low in females, confirming the independent segregation of Chr10 and Chr21. Conversely, LD between Chr10 and Chr21 was very high in males (Figure 3), suggesting that the male haplotypes of those two chromosomes segregate together. One possible mechanism explaining such pattern would be a male-restricted fusion between the Y chromosome and one autosome. Chromosome fission and fusion events are frequent in the evolution of vertebrates, particularly in fish, in which they could explain the enormous species-richness and rapid evolution of karyotypes (Cheng et al 2020; Sutherland et al 2017). Fusions may also be polymorphic in fish and play an adaptive role, as reported in Atlantic salmon (*Salmo salar*), in which fused and unfused karyotypes have been shown to coexist (Lehnert et al 2019; Wellband et al 2019). Moreover, fusions between a Y chromosome and an autosome are known to occur more frequently than autosomal fusions or W-autosome fusions, particularly in fishes Pennell et al (2015). The most probable explanation for the preponderance of Y-autosome fusions involved both selection and sex biases, as the smallest effective size (Ne) of the Y makes it more likely to fix rearrangements including those with partially deleterious effects. This process may be even more prevalent in species with polygynous mating system like fish, which increases the variance of male reproductive success, and in species with skewed sex-ratio, as observed in the halibut. Because of its significant impact on recombination, chromosomal fusion may be one of the key factors underlying the evolution of differentiated sex chromosomes. Fusions are a peculiar class of rearrangements that not only link chromosomes that previously segregated independently but also reduce recombination rates within each arm and prevent recombination between fused and unfused homologous chromosomes in polymorphic populations (Dumas and Britton-Davidian 2002). This typically leads to gradual differentiation from the fusion point as observed here for Greenland Halibut or the young sex chromosomes found in three-spined stickleback (*Gasterosteus aculeatus*) (Kitano et al 2009). This represents a typical step in the birth and establishment of new sex chromosomes (McAllyster 2003, Kirkpatrick 2010), which is often mediated *via* large chromosomal rearrangements such as inversions or fusions. One of the main determinants of this process is selection against recombination, because linkage between sex-determination loci and alleles beneficial to one sex has the potential to resolve sexual conflict (Charlesworth et al 2005). As discussed below, we identified three sex-associated loci spread on two chromosomes and one may speculate whether epistatic interactions may occur between them. Further work would be needed to better understand gene interactions in this region and more formally confirm the putative male-limited fusion (Guerrero and Kirkpatrick 2014; Wright et al 2016).

### Sox2 gene as the major driver of Greenland Halibut SD?

Alignment of DNA or mRNA sequences of previously described sex-associated markers revealed that those genes were widely distributed throughout Greenland Halibut assembly (on seven chromosomes, Table S1). Most of those genes are located in regions of low differentiation between males and females: amhr2, cyp19a1A on chr05; cyp11b2 and rspo1 on chr08; dmrt1 on chr09, foxl2 on chr10 and gsdf on chr18, making them unlikely to underlie sex differentiation. However, three candidate genes mapped to the putative sex chromosomes: gdf6 and *sox2* on Chr10, and *sox9a2* on Chr21 (Figure 2, Table S1), and are thus possible drivers of sex-determination in Greenland Halibut.

Growth differentiation factor 6 (*Gdf6*) is a member of the TGF-β family involved in vertebral segmentation and cell differentiation. While no gonadal function was reported previously, *gdf6Y* emerged recently as a putative SD gene in the Turquoise Killifish (*Nothobranchius furzeri*) (Reichwald et al 2015) because sex-linked allelic variations affect the Gdf6 protein function. The Y chromosome allele, *gdf6Y*, also exhibits sex-linked expression during testicular differentiation in the Turquoise Killifish, supporting its SD role but functional evidence of its role as a SD gene is yet not available.

Sox2 is a member of the SOX (SRY-related HMG-box) family with a high-mobility-group (HMG) DNA-binding domain. It is an important transcription factor, which is involved in developmental processes of vertebrates, such as embryogenesis, maintenance of stem cells, neurogenesis. In the Rohu Carp (*Labeo rohita)*, sox2 mRNA is expressed in various organs, as well as in the culture proliferating spermatogonial stem cells (Patra et al (2015). Sox2 is also a candidate SD gene in two aquaculture species, the mollusc Zhikong scallop (*Chlamys varia*) (Liang et al 2019) and the crustacean black Tiger Shrimp (*Penaeus monodon)* (Guo et al 2019). In flatfish, sox2 is a strong candidate for SD, in Japanese Flounder, sox2 transcripts is expressed in gonadal tissues and the transcript abundance in ovaries is higher than in testis (Gao et al 2014), and, in Turbot, sox2 is defined as a consistent candidate gene putatively driving SD (Martinez et al (2021).

Sox9 gene has been related to sexual differentiation in several teleost species: Rainbow Trout, (*Oncorhynchus mykiss*, Takamatsu et al. 1997; Vizziano et al. 2007), Tilapia (*Oreochromis niloticus*, Ijiri et al 2008), Zebrafish (*Danio rerio* Chiang et al 2001; von Hofsten and Olsson 2005), medaka, *Oryzias latipes* (Yokoi et al 2002), Guppy (*Poecilia reticul*, Shen et al 2007), the Common Carp (*Cyprinus carpio*, Du et al. 2007) and the Asian Swamp Eel (*Monopterus albus*, Zhou et al 2003). In the Rainbow Trout, two sox9 genes have been identified: sox9a1 and sox9a2. Sox9a1 shows higher expression levels in males than in females before sexual differentiation (Vizziano et al 2007). In Tilapia, sox9 expression is similar in XX and XY gonads, but, after sexual differentiation, it is positively regulated in XY gonads (Ijiri et al 2008). In Medaka, testicular sox9 (sox9a2) expression in somatic cells is similar in both sexes during early gonadal differentiation and becomes positively regulated in males during the testicular lobe formation stage (Nakamoto et al. 2005; Nakamura et al. 2008). Overall, these observations support the hypothesis of a significant role for sox9 in either testicular differentiation or testicular development. This gene may also be at play in flatfish since, in Atlantic Halibut, sox9a2 is included in a cascade of gene activation starting with the dmrt1 gene which is activated by the putative SD factor, the gsdf gene (Norris and Carr 2013, Einfeldt et al 2021).

In Greenland Halibut, sox2, gdf6, and sox9a2 represent three possible candidate genes for sex determination. One can think that these three genes may be interacting in shaping the determination of sex and/or the subsequent development of sex organs. However, several lines of evidence suggest that sox2 might have a major role in Greenland Halibut SD. First, sox2 is located in a region that is more highly differentiated between sexes than gdf6 and sox9a2 genes. Furthermore, no functional evidence in sexual determination is available yet, neither for gdf6 nor sox9a2 genes, in other fish. Conversely, sox2 has been reported as the main SD driver for several species including another flatfish (Turbot, Martinez et al 2021).

In contrast with sox2 identified in Turbot, SD genes identified in the two most-closely related flatfish species, Atlantic and Pacific Halibut (Table1), do not appear to differentiate males and females in the Greenland Halibut. This is somewhat unsurprising given that even within the same genus (*Hyppoglossus*), not only different species have different genes driving SD, but they harbor a different heterogametic sex (XY for Atlantic Halibut and ZW for Pacific Halibut, Drinan, Loher, & Hauser 2018, Einfeldt et al 2021).

The nascent SD system with low differentiation between sex chromosomes revealed in Greenland Halibut might reflect the observed complex and still enigmatic sex maturation. For example, in some Greenland Halibut individuals, the maturation cycle can be interrupted by a sudden degeneration of the oocytes (Fedorov, 1971). More recently, it has been suggested that this species utilizes a very unusual oocyte development pattern containing two simultaneously developing cohorts of vitellogenic oocytes, not seen in any related species (Rideout et al 2012; Dominguez-Petit et al 2017). Also, unusual and still unexplained, Greenland Halibut present high densities and regular occurrence of rodlet cells in male and female gonads (Rideout et al 2015). Furthermore, other layers may be added on the top of such an SD mechanism; environmental factors such as temperature, pH, population density, and social interactions have been found to influence the sex-ratio in fish (Crews 1996; Nakamura et al. 1998; Baroiller et al. 1999; Baroiller and D’Cotta 2001). More research would be needed to better disentangle the relative contribution of environmental factors and genetic factors in sex-determination in Greenland Halibut.

## Conclusion

Despite the commercial importance of Greenland Halibut, important gaps still persist in our knowledge of this species, including gaps about its reproduction cycle and sex determination mechanisms. In this study, we used a compendium of novel technologies (single molecule sequencing of long reads (Pacific Sciences) with Chromatin Conformation Capture sequencing (Hi-C) data and whole-genome sequencing of hundreds of individuals) to provide the very first chromosome-level genome reference and elucidate the sex determination mechanism of this demersal fish. The chromosome-level assembly and the investigation of its syntenic relationships with other economically important flatfish species highlight the high conservation of synteny blocks within the flatfish phylogenetic clade. Our sex determination analysis revealed that flatfishes do not escape this rule and exhibit a high plasticity and turnover in sex-determination mechanisms. In fish, the early evolutionary stage of most sex chromosomes and the subsequent small differences between those chromosomes make it extremely difficult to distinguish diagnostic differences between them. A whole-genome sequencing approach of hundreds of individuals allowed us to draw a full picture of the molecular SD system in Greenland Halibut. Specifically, our results suggested that Greenland Halibut has a very nascent male heterogametic XY system, with a putative major effect of the sox2 gene, also described as the main SD driver in the Turbot (*S. maximus*) and to a lesser extent in the Japanese Flounder. Interestingly, our study also suggested for the first time in flatfish a putative Y-autosomal fusion that could be associated with a reduction of recombination typical of early steps of sex chromosome evolution.

## AKNOWLEDGMENTS

We are grateful to Renée Gagné and Léopold Ginther, for assistance while sampling the individual used for the genome assembly and transcriptomic RNAseq and collecting its tissues. We thank all biologists and technicians of Fisheries and Oceans Canada for their implication in the project and their field assistance. We also thank Cécilia Hernandez for help gathering all the samples in the lab and Justine Létourneau for help in DNA extraction. Brian Boyle and Nina Therkildsen provided key advice about sequencing and libraries preparation. The genome assembly was supported by the Canada 150 Sequencing Initiative (Canseq150), CFI and Genome Canada Technology Platform grants to J R. This research was funded by the Canadian Research Chair in Genomics and Conservation of Aquatic Resources, Ressources Aquatiques Québec (RAQ) as well as by a Strategic Project Grant from the Natural Sciences and Engineering Research Council of Canada (NSERC) to L. Bernatchez.

